# Compositional heterogeneity confers selective advantage to protocellular membranes during the origins of cellular life

**DOI:** 10.1101/678847

**Authors:** Susovan Sarkar, Shikha Dagar, Ajay Verma, Sudha Rajamani

## Abstract

Protocells are primitive cellular entities that are thought to have emerged during the dawn of life on Earth. Their membranes are considered to be made up of mixtures of single chain amphiphiles, such as fatty acids and their derivatives, moieties that would have been part of the complex prebiotic chemical landscape. In addition to their composition, the physico-chemical properties of these prebiological membranes would have been significantly affected and regulated by the physical environment that they were present in. In this study, the physico-chemical properties of two different chain length membrane systems were systematically characterized, under pertinent early Earth conditions. The membrane systems were designed to comprise a fatty acid and its alcohol and/or glycerol monoester derivative, to make a range of binary and tertiary vesicle combinations. Their properties were then evaluated as a function of multiple factors including their composition, stability under varying pH, Mg^2+^ ion concentrations and dilution regimes, and their permeability to small molecules. Our results demonstrate how these environmental constraints would have acted as important prebiotic selection pressures to shape the evolution of prebiological membranes. This study also illustrates how different fatty acid derivatives confer varying degree of stability when combined with their respective fatty acid moiety. Interestingly, when the membrane systems were subjected to multiple selection pressures in a consecutive manner, only the heterogeneous membrane systems survived the ‘race’. Our results illustrate that compositionally diverse membrane systems are more stable and robust to multiple selection pressures, thereby making them more suitable for supporting protocellular life.

**Significance statement:** The physico-chemical environment would have played an important role in shaping the composition and evolution of primitive membrane systems. This study demonstrates the importance of compositional heterogeneity on the stability of protomembrane systems under pertinent prebiotic selection pressures. Two different fatty acid based systems were mixed with their respective alcohol and/or glycerol monoester derivatives, to generate combinations of binary and tertiary membrane systems. Increasing chain length effected the physical property of the membranes thus affecting their stability. Furthermore, we also demonstrate how the head groups of the derivatives employed in this study (e.g. glycerol monoester and alcohol) contribute differently towards stabilizing the mixed membranes under a given selection condition, by discerning the molecular mechanism underlying this process. Our results illustrate how multiple selection pressures would have preferentially supported the emergence of compositionally heterogeneous membrane systems.

## Introduction

The earliest forms of cellular life are considered to be entities that comprised of dynamic chemical reactions, encapsulated within amphiphilic compartments (1, 2). Unlike the contemporary biological membranes that potentially resulted over millions of years of evolution, prebiological membranes are thought to have been relatively simpler and composed of single chain amphiphiles (SCAs) (3). These SCAs could have come about on the early Earth either by endogenous synthesis, in the form of Fisher-Tropsch Type (FTT) reactions, or via exogenous delivery (4, 5). In this context, fatty acids and their derivatives have been predominantly studied for their plausible role as early compartments (3, 6). Fatty acids are known to possess high critical vesicular concentrations (CVCs), the concentration at which the monomers assemble into higher ordered structures like vesicles (7). Such high CVC requirement poses significant obstacles towards their self-assembly under prebioitic scenarios, wherein meeting this high concentration prerequisite would have been difficult (8, 9). Furthermore, fatty acid monomer assembly also depends on the pH of the surrounding environment (6, 10). It has been hypothesized that the pH of certain terrestrial hydrothermal pools of the early Earth would have been neutral to alkaline (11). This pH regime is also known to drive prebiotically pertinent reactions, including formose reaction (12), polymerization of non-activated amino acids (13), and non-canonical nucleoside or nucleotide formation (14). Given the sensitivity of fatty acid membranes to changes in pH, the encapsulation of the aforementioned reactions or molecules would have been really challenging. Moreover, fatty acids are also cation sensitive moieties (15). On the contrary, RNA molecules, which are thought to be the first biomolecules to have emerged, require divalent cations in order to efficiently replicate and carry out catalytic functions (16-18). Recent studies have also demonstrated the reconstitution of certain primitive metabolic pathways in the presence of cations (19, 20). The cation concentrations required for facilitating aforesaid reactions have been shown to be incompatible with fatty acid membranes (15, 21) This poses an imminent question of how such metabolic networks and RNA replicators could have been encapsulated in fatty acid based membrane systems.

Nonetheless, fatty acid membranes are dynamic in nature unlike the contemporary phospholipid membranes, and can facilitate the permeation of polar molecules (22). This is an essential requirement for primitive cells as it allows for the exchange of matter with its environment. In this regard, their self-assembly property, permeability and membrane integrity in different environmental conditions, such as pH, ionic strength, temperature, dilution regimes etc., need to be systematically explored. Thus far, studies have predominantly focused on delineating one, or up to two of the aforesaid parameters at any given time (3, 6-9, 21 and 23). However, the aforementioned conditions would have acted in concert as a combination of prebiotic selection pressures, which would have played a vital role in shaping the evolutionary landscape of prebiological membranes. Pertinent to this, it is also rational to envisage the presence of heterogeneous membrane systems as these could have been readily available in an, arguably, large prebiotic chemical space. Previous studies in this regard suggests that the lack of membrane stability can be counterbalanced by increasing the heterogeneity of the membrane system. It is known from literature that addition of long chain alcohols with fatty acids, decreases the CVC of the resultant binary systems (8), and also confers stability to the vesicles at alkaline pH (8, 24). Previous studies have also showed that binary systems of fatty acid and its glycerol monoester are more resistant to soluble monovalent and divalent cations (21, 25). However, the mechanism behind this increase in stability towards divalent cations, in the presence of derivatives, is not clearly understood. Mansy et al. demonstrated that the addition of monomyristolein in myristoleic acid membranes, increases their permeability drastically, whereas addition of myristoleoyl alcohol increases the permeability only slightly when compared to the homogenous myristoleic acid membranes (26). Although insightful, the aforementioned studies predominantly looked at binary membrane systems. Given the heterogeneous nature of the prebiotic soup, and the niche parameters, it would be worthwhile to complexify the starting mix, to better understand how membrane related processes would have advent under ‘prebiotically realistic’ conditions. In this context, a membrane system composed of decanoic acid, decanol and glycerol mono-decanoate, is the only tertiary system that has been explored thus far in terms of its thermostablity and permeability (26, 27).

In order to gain a deeper understanding of how compositional heterogeneity would impinge on a membrane system’s survivability, especially under multiple prebiotic selection pressures, we set out to characterize tertiary membrane systems of selected SCAs. In the present study, fatty acids of two different chain lengths, i.e. oleic acid (OA, C18) and undecylenic acid (UDA, C11), were mixed with their corresponding alcohol and/or glycerol monoester derivatives, and used as a proxy for mixed membrane systems. Fatty alcohols and glycerol monoester derivatives were chosen for further experimentation because of their prebiotic relevance (4, 28). A total of four systems, viz. the homogenous fatty acid system, two binary systems containing fatty acid with either the fatty alcohol or the glycerol monoester derivative, and the tertiary system containing all the three components, were explored for each of the chain lengths. The ratio of fatty acid to its derivative was maintained at 2:1. The prebiotically relevant physical parameters that were characterized include the formation of protocellular membranes at alkaline pH, their CVC, ionic stability and the permeability of the said systems. Our results show that the heterogeneous membrane systems are indeed more stable, and robust under diverse environmental conditions. Therefore, these would have been more suitable to support protocellular life forms. Our results also illustrate that the head groups of the amphiphiles used in this study, play an important role in stabilizing primitive membranes under specific selection conditions. Systems containing different derivatives possess different survival rates when subjected to a specific selection pressure. Given this interesting finding, we also attempted to delineate the contribution of individual head-groups, and the plausible mechanism that might be involved in stabilizing the protomembrane systems that were evaluated. Furthermore, the possibility of being simultaneously affected by multiple selection pressures, is a prebiotically “realistic” scenario. An important aspect of our study has been to understand this pertinent scenario and results from this aspect of our study clearly demonstrates that the tertiary systems would have possessed the best chance at survival when subjected to multiple selection pressures. The overall outcome of this work illustrates that the evolution of protomembranes would have been shaped, both, by their compositional heterogeneity, and the niche parameters (selection conditions) that these systems were subjected to. This work, thus, has implications for discerning the emergence of mixed membrane systems, and highlights the need to factor prebiotically realistic conditions, to better understand how they would have impinged on the evolutionary landscape of prebiotic membranes.

## Results

### Influence of compositional heterogeneity on self-assembly of protocellular membrane systems

All the experiments described in this present study were conducted using binary and tertiary mixed membrane systems of C11 and C18 fatty acids. The binary systems based on the undecylenic acid (C11) fatty acid system were prepared by mixing undecylenic acid (UDA) with either glyceryl 1-undecylenate (UDG) or undecylenyl alcohol (UDOH) in 2:1 ratio. The tertiary system was prepared by mixing all three components together, i.e. UDA:UDG:UDOH, in a ratio of 4:1:1. Similarly, the oleic acid based (C18) binary systems were prepared by mixing oleic acid (OA) either with glycerol 1-monooleate (GMO), or the oleyl alcohol (OOH) derivative in a ratio of 2:1, and the tertiary system was made by mixing all the three components together (OA:OOH:GMO system) in 4:1:1 ratio. Figure S1 (SI) summarizes the structures of the aforesaid amphiphiles used in this study. In order to determine the CVC of the membrane systems, 1,6-diphenyl-1,3,5-hexatriene (DPH) was used as a bilayer reporter fluorescent probe (SI, Method section). DPH fluorescence was plotted as a function of lipid concentration. The concentration of lipid where a sudden increase in fluorescence is observed (the inflection point) represents the CVC of the system (29). Additionally, turbidity assay and microscopy were also performed to confirm the CVC results (SI, Figure S4) (7). The fluorescence assay revealed that the CVC of the homogenous UDA was near about 35 mM. Upon mixing UDA with UDG, the CVC drastically decreased to 2 mM (SI, Figure S2). For the binary system of UDA:UDOH and the tertiary system of UDA:UDG:UDOH, the CVC was found to be around 3 and 2 mM, respectively (SI, Figure S2). Microscopic analysis of all the systems confirmed the presence of vesicles (SI, Figure S4). For the homogenous oleic acid (OA) system, the CVC was found to be around 0.09 mM. Upon adding either the GMO or the OOH derivative, the CVC decreased to about 0.02 and 0.06 mM, respectively (SI, Figure S3). For the tertiary system of OA:GMO:OOH, the CVC was found to be around 0.02 mM (SI, Figure S3). On performing microscopy, vesicles were observed under the microscope at 40X magnification only at a slightly higher concentration than what was expected from the fluorescence assay (as is summarized in Table T1). This could potentially stem from the fact that light microscopy is diffraction-limited thus missing out on vesicles that are smaller than 200 nm.

**Table T1:** Summary of the CVCs of all the different C18 and C11 systems using three different assays. Columns 2, 3, and 4 provide a comparison of the difference in the CVC estimation using fluorescence assay, microscopy and turbidity assay, respectively. The molar ratio of fatty acid to overall derivative was kept to 2:1. UDA, undecylenic acid; UDG, glyceryl 1-undecylenate; UDOH, undecylenyl alcohol; OA, oleic acid; GMO, glycerol 1-monooleate; OOH, oleyl alcohol.

| System used | Apparent CVC in mM |  |  |
| --- | --- | --- | --- |
|  | Fluorescence Assay | Microscopy | Turbidity Assay |
| UDA | 35 | 42 | 40 |
| UDA:UDOH | 4 | 12 | 7 |
| UDA:UDG | 2 | 2 | 3 |
| UDA:UDG:UDOH | 2 | 2 | 2.5 |
| OA | 0.09 | 0.6 | 0.1 |
| OA:OOH | 0.06 | 0.1 | 0.1 |
| OA:GMO | 0.01 | 0.1 | 0.1 |
| OA:GMO:OOH | 0.02 | 0.05 | 0.08 |

### Effect of compositional heterogeneity on the formation of protocellular membranes under alkaline pH regimes

Effect of compositional heterogeneity on the formation and stability of the protocell membranes was investigated from pH 7 to 11 (SI, Method section). It was observed that both the alcohol and glycerol monoester derivatives could indeed stabilize the fatty acid vesicles over a wide range of pH. Turbidity estimation indicated that UDA alone could form vesicles from pH 7.5 to 8 (SI, Figure S7), which was also confirmed using microscopy (SI, Figure S5). The binary UDA:UDG system formed vesicles from pH 7 to 9, above which micelle formation resulted in a decrease in the turbidity values (SI, Figure S7). On the other hand, the UDA:UDOH binary system and the tertiary system of UDA:UDG:UDOH were able to form vesicles, all the way from pH 7.5 to 11 (SI, Figure S5). As for the OA based systems, the homogenous OA system formed vesicles from pH 8 to 9 (SI, Figure S6). Below pH 8 and above 9, disordered aggregates and micelles, respectively, were observed. The OA:OOH binary system could form vesicles from pH 8.5 to 11. However, below 8.5 it assembled into disordered aggregates (SI, Figure S6). In the case of the OA:GMO binary system and the tertiary system of OA:GMO:OOH, vesicles were observed over a wide range of pH starting from 7.5 and up to 11. Below pH 7.5, both the systems only formed aggregates. The aforesaid results have been summarized in Figure 1.

**Figure 1:**
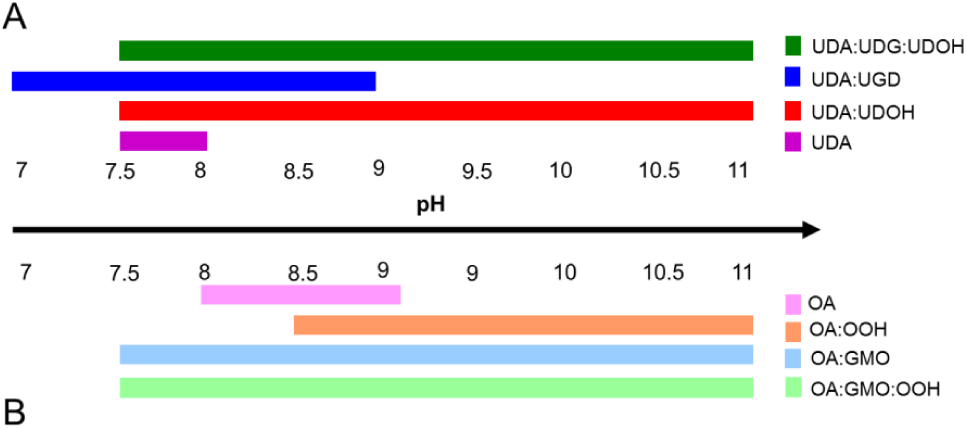
Illustration showing the ability of the different fatty acid-based systems to form vesicles over varying pH regimes. The ability of a system to form vesicles over a range of pH is represented by the differently colored horizontal bars. The length of the bar is directly proportional to the range of pH at which the system can assemble into vesicles. Panels A and B represent the UDA and OA based mixed systems, respectively. The molar ratio of fatty acid to overall derivative was kept to 2:1.

### Effect of membrane heterogeneity on its stability in the presence of Mg^2+^ ions

Previous studies have highlighted the role of Mg^2+^ ions on the function of ribozymes (16-18). In this aspect of the work, we aimed to understand the influence of alcohol and glycerol monoester head groups, towards the ionic stability of fatty acid in the presence ofMg^2+^ ions. Optical microscopy and dynamic light scattering (DLS) spectroscopy were used to check the vesicle stability (SI, Method section). The concentration of Mg^2+^ ion at which the average size of the vesicle population increases, in comparison to the initial size (when no Mg^2+^ is added), was considered to be the aggregation-inducing concentration (Mg^2+^AIC) (30). At this concentration, both, vesicles and Mg^2+^ ion induced aggregates can be present simultaneously in the solution. DLS analysis showed that the UDA:UDG binary system was found to be the most stable one, with an Mg^2+^AIC of 16 mM, followed by the tertiary UDA:UDG:UDOH and the binary UDA:UDOH systems, with an Mg^2+^AIC of 14 and 8 mM, respectively (SI, Figure S8). The UDA system was extremely labile to Mg^2+^ ions, where aggregation started even at Mg^2+^ ion concentrations of as low as 3 mM of (SI, Figure S8). On microscopic analysis, large fatty acid crystals and aggregates were observed in the UDA system at 4 mM Mg^2+^ ion concentration, and no vesicles were observed in the solution beyond 8 mM Mg^2+^ concentration (Figure 2, Panel A). However, in the case of the UDA:UDG system, lipid aggregates and crystals started appearing at 12 mM Mg^2+^ ion concentration, and vesicles persisted even in solutions containing a Mg^2+^ ion concentration of 24 mM (Figure 2, Panel B). Between the UDA:UDOH and the UDA:UDG:UDOH systems, the latter seemed more stable, with vesicles being observed along with some aggregates in the presence of 12 mM Mg^2+^ ions (Figure 2, Panel D). However, only crystals and droplets were found in the UDA:UDOH system at 12 mM Mg^2+^ ion concentration under the microscope (Figure 2, Panel C). Among the four C18 based systems, the OA alone system was found to be the most sensitive, with an Mg^2+^AIC of 3.5 mM (SI, Figure S9). Interestingly the OA:GMO was found to be the second most sensitive towards Mg^2+^ ions, with an Mg^2+^AIC of 5 mM. Both, the binary OA:OOH and the tertiary OA:GMO:OOH systems showed an Mg^2+^AIC of 6 mM (using DLS) (SI, Figure S9). Similar observations were confirmed using microscopy (SI, Figure S10). Table TS1 in the supplementary information compares Mg^2+^AIC of all eight membrane systems across the two experimental methods used. Overall, the Mg^2+^AIC of all four C18 based systems were lower compared to the C11 systems because total of 2 mM lipid concentration were used for all four C18 systems, which is much lower that were used for C11 based systems.

**Figure 2:**
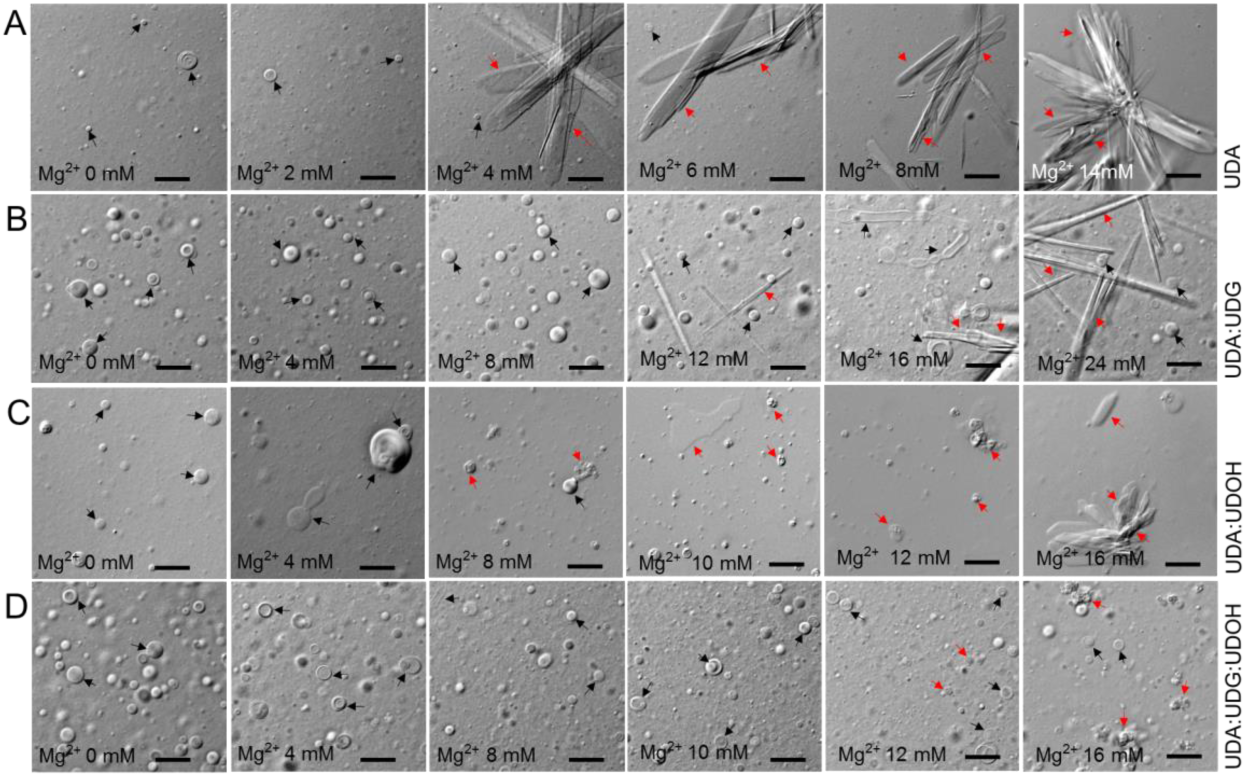
Mg^2+^ ion induced vesicle and aggregate forming properties of all the four UDA based systems (panels A to D). From left to right, Mg^2+^ ions concentration was increased gradually by keeping the lipid concentration constant. In terms of cation sensitivity among the four systems, the following order is seen: UDA > UDA:UDOH > UDA:UDG:UDOH > UDA:UDG. The black and red arrows indicate vesicles and aggregates (fatty acid crystals and droplets), respectively. The scale bar in all the images is 10 microns.

### Influence of compositional heterogeneity on the permeability of protocell membranes

In order to understand the effect that each head group might have on the permeability of model protocell membranes, calcein leakage assay was carried out (SI, Method section). Calcein is a small molecule with a size comparable to that of a dinucleotide, and it possesses three negative charges at the pH 8. C11 based systems were chosen to explore this aspect in detail because of their intrinsic dynamicity coming from them having a smaller chain length. Previous studies have looked at the permeability of homogenous OA and binary OA:GMO based system (25, 26, 31). However, we are not aware of any other study that has looked at the permeability of any C11 based systems. In a typical experiment, calcein was encapsulated in the different vesicular systems above its self-quenching concentration, and its release was measured over time to estimate permeability of the membrane systems, which is in turn a reflection of its dynamicity (30).

It was observed that when UDG was mixed with UDA, the permeability of this mixed binary membrane system increased drastically, in comparison to the pure UDA system. Over a period of 180 min, 80 percent of the encapsulated calcein was released from the UDA:UDG system (Figure 3A, blue trace), whereas 22 percent of the encapsulated calcein was released in case of the UDA membrane system (Figure 3A, purple trace). Interestingly, the UDA:UDOH binary system was found to be impermeable to calcein, where none of the encapsulated calcein was released even after a period of 180 min (Figure 3A, red trace). The tertiary system (UDA:UDG:UDOH) was found to possess moderate permeability to calcein. It was less permeable than the binary UDA:UDG binary system and the homogenous UDA system, but more permeable than the UDA:UDOH binary system, releasing only 14 percent of encapsulated calcein over 180 mins (Figure 3A, green trace). While presence of UDG in the membrane led to an increase in the permeability, the presence of UDOH decreased the permeability of the UDA system. Additionally, the presence of both of the derivatives resulted in an intermediate trait.

**Figure 3:**
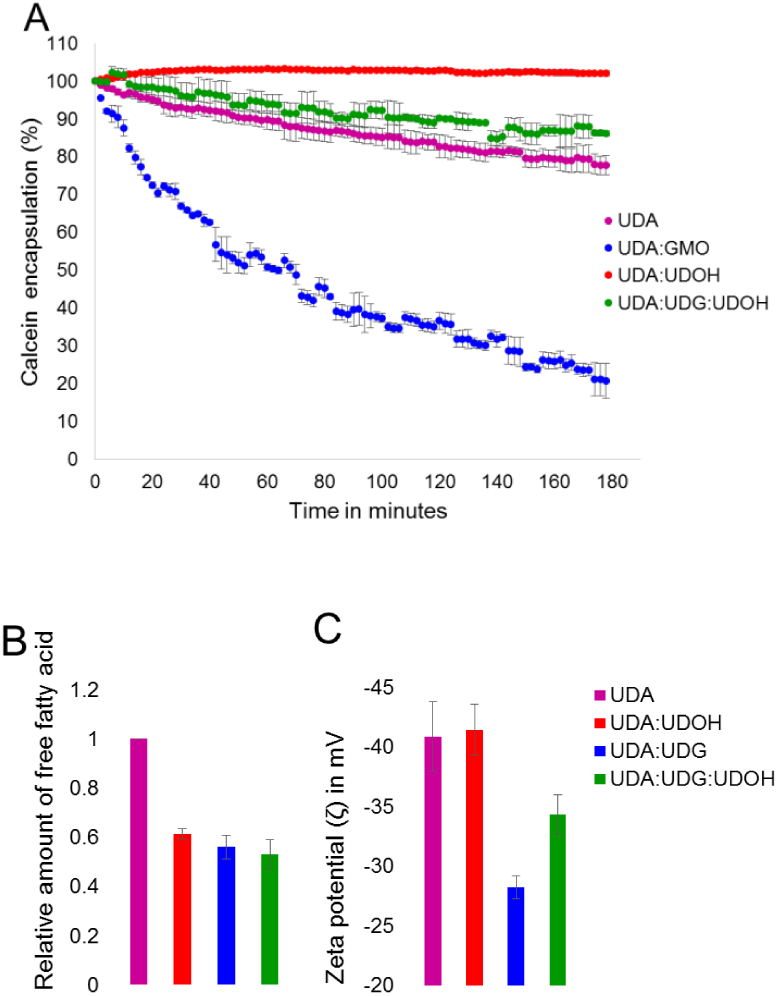
3A. Calcein leakage assay to evaluate the permeability of the four different UDA based system towards calcein. Percentage of calcein encapsulation was plotted against time (in minutes). Decrease in the % encapsulation, indicates increased permeability. n = 4; error bars represent standard deviation (s.d.). 3B. Shows the relative amount of free fatty acids in the solution as a function of the membrane composition. Presence of the derivatives UDG and UDOH increases the retention of UDA in the membrane. n = 6; error bars represent standard deviation (s.d.). The difference between the mean values for the homogenous UDA system with other three heterogeneous systems is significant based on one-tailed student t test with a *p*-value < 0.05. 3C. Indicates the zeta potential measurements of the various C11 systems as a function of their composition. n = 5; error bars represent standard deviation (s.d.). The difference between the mean values for UDA and UDA:UDG is significant based on one-tailed student t test with a *p*-value < 0.00001. The difference between the means for UDA and UDA:UDG:UDOH is significant based on one-tailed student t test with a *p*-value < 0.00083.

### Mechanism by which compositional heterogeneity might confer stability on fatty acid membranes in the presence of Mg^2+^ ions

In order to understand the mechanism behind the increased stability of fatty acid membranes towards Mg^2+^ ions in the presence of alcohol and glycerol monoester derivatives, the charge density associated with the membrane systems, and the retention of the fatty acid moieties in the membranes, was investigated. Fatty acid molecules stay in dynamic equilibrium and can therefore readily exchange between the membrane phase and the free monomers present in the solution (32). The free monomers possess a negative charge and can interact with the free Mg^2+^ ions, potentially forming crystals and unordered aggregates. Therefore, more the amount of free fatty acids in the solution, greater will be the aggregate formation. It is known that phospholipids can increase the retention of fatty acid molecules in the membrane, resulting in a decrease in the free fatty acids, thus stabilizing the membrane in the presence of Mg^2+^ ions (33, 34). In this study, the free fatty acid monomers present in the solution was estimated for all the eight membrane systems using liquid chromatography coupled with mass spectrometry (SI, Method section). The amount of free fatty acid for each of the heterogeneous systems was normalised to their respective homogenous fatty acid system. It was observed that the UDA system possessed the highest amount of free fatty acids (Figure 3B, purple bar). The presence of UDG and UDOH showed a decrease in the dissociation of free UDA molecules into the solution, thereby stabilizing the mixed membranes (Figure 3B, blue and red bar). Nonetheless, there was no significant difference in the amount of free fatty acids between the three heterogeneous systems. In case of OA based membrane systems, it was observed that both, OOH and GMO decrease the dissociation of OA into the solution, similar to what was observed in the UDA based systems (SI, Figure S12, Panel A).

The surface charge density of the membranes was estimated by measuring their zeta potential (ζ) (SI, Method section). The free Mg^2+^ ions present in the solution also tend to interact with the negatively charged fatty acid vesicles and initiate aggregation. Therefore, a decrease in the net negative charge on the vesicle surface would lead to a weakening in the interaction with the Mg^2+^ ions. It has been previously argued that presence of, both, long chain alcohols and glycerol monoesters in fatty acid membranes, can decrease the negative charge density on the membrane (26). Dalai et al. showed that the addition of phospholipids to oleic acid membranes can decrease the zeta potential of the membranes, which in turn stabilizes the fatty acid membranes in the presence of Mg^2+^ ions (30). The zeta potential measurements revealed that the addition of UDG to the UDA membrane, decreased the net negative charge density significantly (Figure 3C, blue bar). The net negative charge of the UDA:UDOH binary system was found to be equal to that of the only UDA system (Figure 3C, purple and red bar) The negative charge on the tertiary UDA:UDG:UDOH system was found to be somewhere in between these, i.e higher than the UDA:UDG system but lower than the UDA and UDA:UDOH system (Figure 3C, green bar). The change in the negative charge density can be explained by considering the size of the different head groups. Because of the bulky head group of the UDG moiety, less number of deprotonated fatty acid molecules would be present in a given unit surface area. This is in contrast to the number of deprotonated fatty acid molecules that could be present in the same unit surface area of the UDA and the UDA:UDOH membrane systems. This observation is further strengthened when taking into account the permeability of the membrane systems as shown in Figure 3A. The presence of bigger head group leads to looser packing, thus increasing the overall permeability of the said system. Similar pattern of zeta potential values was also observed for the oleic acid based membranes. The OA:GMO binary system was found to possess the lowest negative charge (SI, Figure 12, Panel B, blue bar). Whereas the OA:OOH and OA membranes contain similar negative charge densities on them (SI, Figure 12, Panel B, purple and orange bar). Therefore, we found that both of the components, i.e. retention of fatty acid molecules in the membrane, and the change in the surface charge density, would contribute towards the increased stability of the mixed membrane systems in the presence of Mg^2+^ ions. Long chain alcohols stabilize the fatty acid membranes by retaining the fatty acid molecules in the membrane, whereas the glycerol monoester not only increases the fatty acid retention in membrane, but also decreases the overall negative charge density.

### Multiple selection pressures (MSPs) and survivability of protocell membranes as a function of their composition

The stability of different membrane systems under multiple selection pressures (MSP) was investigated next using C11 based systems. The three selection pressures discussed above i.e. dilution regimes, formation of vesicles in alkaline pH and stability in the presence of Mg^2+^ ions, were applied in a sequential manner. Furthermore, to understand if there was any influence coming from the order of the applied selection pressures, a total of six different combinations of the aforesaid selection pressures were applied by varying their sequence. After applying each selection pressure, the solution was observed under microscope to check for the presence of vesicles (SI, Method section). The MSP experiment revealed that even though the binary mixed systems (UDA:UDOH and UDA:UDG) could be more stable under a given selection pressure, when all the three selection pressures were applied sequentially, the tertiary system (UDA:UDG:UDOH) was the one that stood the best chance at survival. As shown in Figure 4 panel A, all four C11 based systems formed vesicles at a lipid concentration of 60 mM at pH 8. When all the four systems were diluted to a final lipid concentration of 20 mM using a buffer of pH 8, the UDA system failed to form vesicles because of its high CVC (near 35 mM), while the other three mixed systems continued to form vesicles (Panel B). Next, out of the three mixed systems, only lipid crystals and aggregates were observed in UDA:UDOH system when Mg^2+^ ions were added to the solution at a concentration of 14 mM (Panel C). Among the other two mixed systems, UDA:UDG and UDA:UDG:UDOH, being less sensitive to Mg^2+^ ion concentration, continued to form vesicles. Finally, when the pH of the solution was adjusted to 10, the UDA:UDG system failed to assemble into vesicles. Consequently, only the tertiary system of UDA:UDG:UDOH was able to assemble into vesicles when all the selection pressures were applied back to back (Panel D). Significantly, this observation was independent of the sequence of the selection pressures applied (SI, Figure 13 to 17).

**Figure 4:**
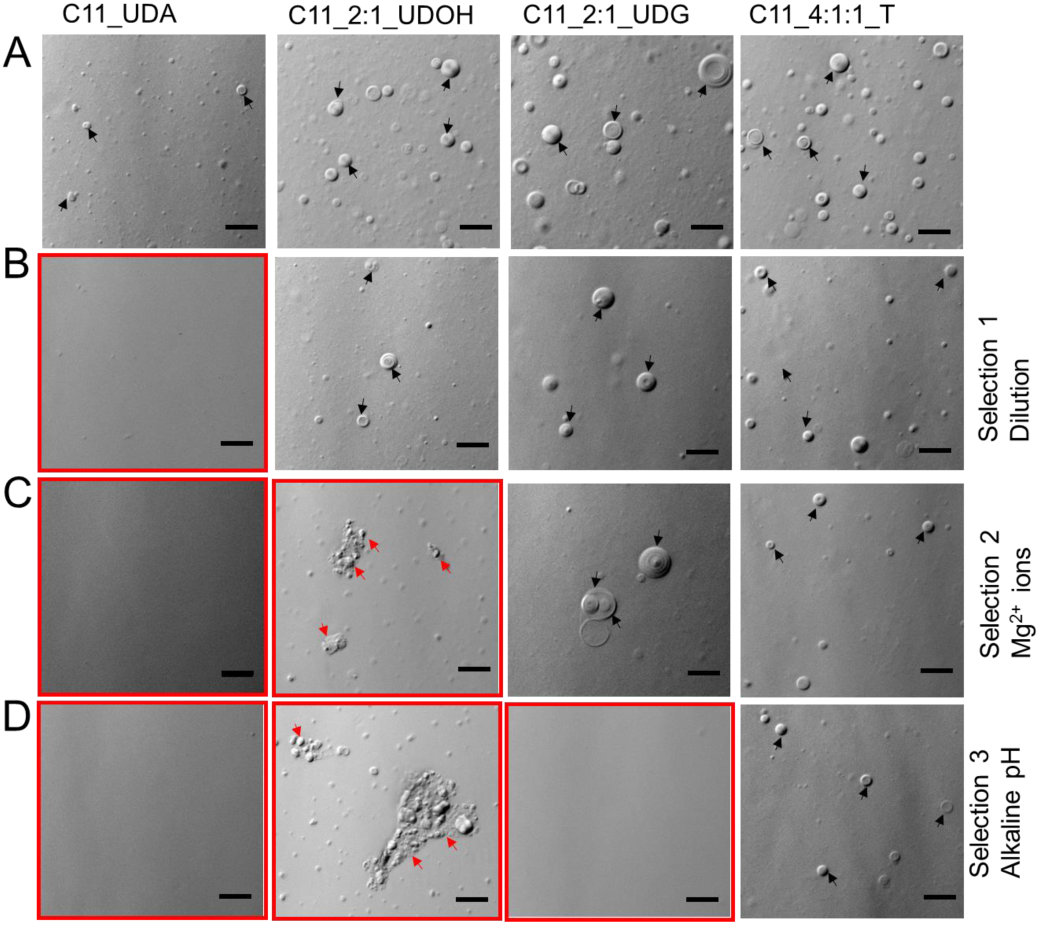
Vesicle stability as a function of their composition under multiple selection pressures when applied sequentially. Panels B to D represent different pebiotically relevant selection conditions. A) All four C11 systems at a concentration of 60 mM at pH 8. B) Dilution regime selection: All four C11 systems diluted to a concentration of 20 mM at pH 8. C) Stability in presence of Mg^2+^ ions: All four C11 systems at 20 mM total lipid concentration at pH 8, in the presence of 14 mM Mg^2+^. D) Stability at alkaline pH: All systems at 20 mM total lipid concentration in the presence of 14 mM Mg^2+^ and at an alkaline pH of 10. The red enclosures indicate absence of vesicles. The black and red arrows indicate vesicles and aggregates (fatty acid crystals and droplets), respectively. The scale bar in all the images are 10 microns.

## Discussion

It is reasonable to assume that the environments of the early Earth would have been heterogeneous and replete with different kinds of amphiphiles that could readily assemble into membrane structures under pertinent conditions. The physico-chemical properties of these primitive compartments would have been largely affected by their environmental conditions. In the present study, the formation and stability of model protomembrane systems, and the influence of membrane compositional heterogeneity on their fitness, was systematically evaluated under different early Earth selection conditions. When the fitness of the different membrane systems were tested using dilution as the selection regime, it was observed that the incorporation of both of the derivatives, i.e. fatty alcohol and glycerol monoester moieties, decreased the CVC of the system. The UDA:UDG and the UDA:UDG:UDOH systems were found to possess the lowest CVC among the four UDA based systems (Table 1). While, in the OA based systems, the OA:GMO system was found to have the lowest CVC of about 0.01 mM, which suggests that the influence of the glycerol monoester on lowering the CVC is possibly greater than the fatty alcohol moiety. A crucial aspect to highlight here is that three different methods were employed in our study to estimate the CVC. This is because use of any of the individual techniques in isolation is not sufficient to gain an understanding of the complete picture. For e.g., DPH can bind to, both, the higher order amphiphilic structures, i.e the ordered vesicles, and to the disordered random aggregates (29). As for CVC estimation by turbidity measurement, both, vesicles and aggregates can contribute to the resultant turbidity of a solution, giving false positive values, while the microscopic analysis tends to miss out on smaller sized vesicles due to diffraction limitation (7, 35). Given the aforesaid limitations of the individual techniques, the three techniques were combined to confidently narrow down the CVC of the systems to a precise range.

As for the stability of the various systems in alkaline pH regimes, homogenous fatty acid systems were found to be extremely sensitive (SI, Figure 5, Panel A). The UDA:UDG system failed to form vesicles above a pH of 9. Whereas, the UDA:UDOH binary system and the UDA:UDG:UDOH tertiary system continued to assemble into vesicles even at pH 11 (SI, Figure 5). In case of the OA based systems, all three mixed systems (OA:GMO, OA:OOH, OA:GMO:OOH) were found to form vesicles even at pH 11 (SI, Figure 6). These observations can be explained by factoring in the protonation status of the different species. As the pH increases, the fatty acid species get deprotonated, allowing it to hydrogen bond with the hydroxyl group of the alcohol or the glycerol head group, resulting in membrane assembly (3, 6). Hence in case of UDA:UDG:UDOH tertiary system, the membrane stability predominantly seems to stem from the UDOH moiety above pH 9 as the UDA:UDG system fails to assemble into vesicles above pH 9. As for the OA:OOH system, it does not form vesicles at pH 8, while the pure OA system can (SI, Figure 6, Panel C). This might potentially come from the predominance of the protonated species at this pH, which hampers efficient hydrogen bonding. The ability of the OA:GMO and UDA:UDG binary systems, to assemble into vesicles at lower pH, while the corresponding homogenous fatty acid systems could not, indicates that at the lower pH regimes, the stability of the mixed membrane is predominantly conferred by the glycerol monoester moiety. These observations indicate to a pH range within which the derivative in the membrane could positively influence membrane stability.

When the stability of the membrane systems in question was evaluated against Mg^2+^ ion concentration, the UDA:UDG binary system was found to be the most resilient among the UDA based systems, followed by the UDA:UDG:UDOH (SI, Table TS1). The UDA:UDOH was found to be the second most sensitive one after the homogenous UDA system. In the OA based systems, the OA:OOH and the OA:GMO:OOH systems were found to be equally stable towards higher Mg^2+^ ion concentrations. The OA:GMO binary system was found to be the second most sensitive one, after the only OA-based system (SI, Table TS1). The grater stablyzing effect of OOH than that of GMO could potentially be attributed to the increase in the chain length. As the chain length increases, parameters like membrane thickness, membrane packing, CVC etc. also change. Previous studies have demonstrated that the presence of chelating agents such as citrate could stabilize oleic acid membranes in the presence of high concentrations (50 mM) of Mg^2+^ ion, while allowing for the function of the encapsulated ribozyme. (36, 37). However, presence of chelated Mg^2+^ decreases the rate of the reaction significantly (34, 36). This becomes very relevant when low amount of Mg^2+^ ions are present in the environment, highlighting the disadvantage of chelated Mg^2+^ ions (34). Meeting the high concentration requirement of Mg^2+^ ions, and the even higher concentrations of certain chelating agents, might have been difficult in the dilute regimes of the prebiotic Earth. It has been reported that, low concentration of free Mg^2+^ ions (4 to 15 mM) can facilitate ribozyme function (21, 25, and 38), which are compatible with the mixed membrane systems described in this study. Therefore, it seems reasonable to hypothesize that an increase in the membrane compositional heterogeneity, along with some chelation, could have provided a respite from the Mg^2+^ ion conundrum.

Lacking a complex transport machinery, protocells would have depended on membrane dynamics for the transport of nutrients and waste across its boundary (22). Recent studies suggest potential pathways for the transition from fatty acid to phospholipid rich membranes (30, 34). However, it is well known that the incorporation of phospholipids into fatty acid membrane drastically decreases their permeability by decreasing the membrane dynamics (22, 30, and 34). Compartments possessing very high or very low permeability would not have been suitable to support protocellular life forms. Rather, compartments possessing moderate permeability, which would have allowed the permeability of small molecules to the inside, without letting the internal components to permeate out, would have been more ideal. Results from our permeability experiments (Figure 3A) show that the incorporation of UDG into UDA membranes in 2:1 ratio, increases the membrane permeability when compared to the homogenous UDA membrane system. On the other hand, the addition of UDOH decreases the permeability of the system.

Therefore, an optimally permeable membrane would have required a mixture of both of these derivatives, as observed in the tertiary system of UDA:UDG:UDOH. Furthermore, because of the bulky head group, the glycerol monoester is thought to hinder efficient membrane packing, while also stabilizing membrane curvatures, which might have resulted from membrane solute interaction (26, 39). Thus, the permeability of the system increases. UDOH on the other hand, possess a small head group, which can potentially increase membrane packing, resulting in a decrease in the permeability. Basically, tuning the concentration of the various components could, in turn, enable the tuning of the permeability of a given protocellular system that would have determined its suitability for a specific set of environmental parameters. A recent study from Rendón et al. demonstrated that in the presence of fatty alcohol, the vibrational freedom of the carboxyl group of the fatty acid decreases, as a result of increased packing, further supporting our aforementioned hypothesis (24).

In conclusion, our study highlights that the stability of model protocellular membranes is not a linear property of its compositional heterogeneity. This is because, it is governed differently by different prebiotically relevant parameters. Importantly, the effect of each head group on the stability of the membrane, depends on the environment and the selection pressure that the system is being subjected to. For example, at alkaline pH of 10 or above, the UDOH stabilizes UDA membranes, while UDG cannot. Whereas, in the presence of Mg^2+^ ions, the UDG stabilizes the UDA membranes more than the UDOH moiety. Overall, the tertiary system was found to be more stable in the presence of all the three selection conditions, when they were applied sequentially. When put together, these results indicate that different prebiotically pertinent selection pressures would have shaped the evolution of protocellular membranes in a specific manner that was predominantly determined by their composition. Significantly, when multiple selection conditions act upon concurrently, the tertiary systems, being more complex, possess a better chance at survival, highlighting that complex mixed membrane systems would have been necessary to support the emergence of protocellular life forms on the early Earth.

## Materials Methods

Materials and Methods are discussed in detail in the supplementary information (SI).

## Acknowledgments

The authors wish to acknowledge the Microscopy facility and the Mass spectrometry facility (co-funded by DST-FIST and IISER Pune) at IISER Pune. SR acknowledges Department of Biotechnology, Govt. of India [BT/PR19201/BRB/10/1532/2016] for extramural funding. SS acknowledges the research fellowship received from IISER Pune. SD acknowledges the research fellowship received from CSIR, Govt. of India.

## Author contributions

S.S. and S.R. designed research; S.S., S.D and A.V. performed research; S.S., S.D., and S.R. analyzed data; S.S. and S.R. wrote the paper.

