## Supplementary File for "Compositional heterogeneity confers selective advantage to protocellular membranes during the origins of cellular life"

**Content:**

Materials

Methods

Table TS1

Figures S1 to S17

**Materials**

**Materials**

Magnesium chloride hexahydrate ($MgCl_2 \cdot 6\ H_2O$, 203.30 g/mol), sodium hydroxide (NaOH, 39.997 g/mol), hydrochloric acid (HCl, 37 %, 36.46 g/mol), bicine ($C_6H_{13}NO_4$, 163.17 g/mol), CHES ($C_8H_{17}NO_3S$, 207.287 g/mol), calcein (622.55 g/mol) and triton X100 (647 g/mol) were purchased from Sigma Aldrich (Bangalore, India) and used without further purification. All the fatty acids mentioned in this study, namely oleic acid (cis-9, $C_{18}H_{34}O_2$, 282.47 g/mol), oleyl alcohol (cis-9, $C_{18}H_{36}O$, 268.478 g/mol), glycerol 1-monooleate (cis-9, $C_{11}H_{40}O_4$ 356.547 g/mol)**,** undecylenic acid ($C_{11}H_{20}O_2$, 184.279 g/mol), undecylenyl alcohol ($C_{11}H_{22}O_1$**,** 170.29), glyceryl 1-undecylenate ($C_{14}H_{26}O_4$**,** 258.35 g/mol) and myristoleic acid (226.36 g/mol) were purchased from Nu-Chek-Prep (Elysian, MN, USA) and used without further purification. All other chemicals were purchased from Sigma Aldrich (Bangalore, India) and used without further purification. All the experiments were carried out using Nanopure (18 MΩ-cm) water.

**Methods**

**Vesicle solution preparation**

The vesicle solutions were prepared by dissolving the desired amount of the fatty acid and its derivatives in chloroform at a concentration of 10 mg/ml. The chloroform solution was dried under nitrogen gas flow to prepare a dry lipid film. It was then kept under vacuum for five to six hours to make sure that no trace amount of chloroform remained. Subsequently, different buffers (bicine or CHES) of desired pH were used to rehydrate the thin film to form the vesicles. This vesicle suspension was heated for one hour at 60 °C to maximize vesicle formation.

**Microscopic analysis**

Lipid samples were observed under 20X and 40X magnification using a Differential Interference Contrast (DIC) microscope AxioImager Z1 (Carl Zeiss, Germany), (NA = 0.75) to observe the presence of different higher order aggregates. Typically, 10 µL of lipid solution was spread on a glass slide, followed by placing an 18X18 mm coverslip on top of it and covering the four sides with liquid paraffin to decrease the motion of the lipid solution. Thereafter, the slide was immediately observed under the microscope.

**Estimation of critical vesicular concentrations (CVCs) of the membrane systems**

Three different methods were used to estimate the CVC of the membrane systems in question. The first method involved using 1,6-diphenyl-1,3,5-hexatriene (DPH), a hydrophobic fluorescent dye, that can partition into the hydrophobic region of the membrane. Upon partitioning, its fluorescence increases several folds. The increase in fluorescence is directly proportional to the lipid concentration. DPH fluorescence also depends on the kind of lipid used, amount of dissolved ions, and incubation time. The lipid solution used in the experiment was prepared by diluting a stock lipid solution with a pH appropriate buffer. For the C11 and C18 systems, 200 mM bicine buffer of pH 8, and 100 mM CHES buffer of pH 9, respectively, was used to rehydrate the dried lipid film and to prepare further dilutions. The lipid suspension was then sonicated to form a homogeneous mixture of small unilamellar vesicles. In a typical reaction, 1.8 µL of 400 µM methanol solution of DPH, was added into 180 µL of C11 lipid solution to achieve 4 µM final concentration of DPH in the solution. For all the C18 systems the final DPH concentration was decreased to 2 µM to account for reduced lipid concentration (long chain fatty acids tend to have much lower CVCs than small chain fatty acids). The lipid solutions were prepared by diluting the stock lipid solution with appropriate buffer. After adding the DPH into the lipid solution, the mixture was kept at 40° C at a constant rotation of 700 rpm for 30 minutes to increase the partitioning of DPH in to the membrane. Post-incubation, the solution was transferred to a 96-well plate. The fluorescence was measured using a 96-well plate reader on Thermo Scientific Varioskan Flash multimode reader (Thermo Scientific, Singapore) by exciting the samples at 350 nm and measuring the emitted light at 452 nm. To corroborate the CVC values obtained from this fluorescence assay, the turbidity of the lipid solution was also measured at 400 nm, which is a widely used technique in the field, for reporting CVCs. The lipid concentration at which the turbidity of the system increases sharply is considered to be an indication of formation of vesicles, and hence is considered as the CVC of the system. A UV-1800 UV-Vis Spectrophotometer (Shimadzu Scientific Instruments Inc., Columbia, USA) was used to check the turbidity of the lipid solutions, which was indicative of higher order structure formation. The presence of vesicles was further confirmed by microscopy at 40X magnification, as described in the aforementioned section.

**Evaluation of formation of protocellular membranes in alkaline pH regimes**

The ability of different lipid systems to assemble into vesicles was evaluated from pH 7 to 11, at intervals of 0.5 pH units (e.g. pH 7, 7.5, 8 and so on). 6 and 60 mM lipid concentration was used for the C18 and C11 systems, respectively. Typically, the dried lipid films were hydrated with a buffer of appropriate pH so as to cover the whole pH range mentioned. This was done considering the fact that different buffers have their own range of buffering capacity. For example, 200 mM bicine was used to prepare buffers in the pH 7 to 9 regime, while 200 mM CHES was used for pH 9.5 to 11 regime. The scattering of the solution at 400 nm was used as a proxy to gauge the presence of higher order lipid assemblies that were present in the solution using UV-1800 UV-Vis Spectrophotometer (Shimadzu Scientific Instruments Inc., Columbia, USA). The same samples were subsequently observed under microscope at 40X magnification to discern the nature of the higher order assemblies (e.g. vesicles, droplets etc).

### Stability of vesicles in the presence of $Mg^{2+}$ ions

In order to check for the stability of the vesicles in the presence of $Mg^{2+}$ ion, Dynamic Light Scattering (DLS) spectroscopy method was used. In a typical experiment, the vesicle suspension was extruded 15 times through a 200 nm size cut-off polycarbonate membrane using Avanti mini extruder (Avanti Polar Lipids Inc., Alabaster, AL, USA). The $Mg^{2+}$ ions were then added to the lipid solution by adding a desired volume of $MgCl_2$ stock solution, prepared in the respective buffer. 100 mM CHES buffer of pH 9 and 200 mM bicine buffer of pH 8 were used to prepare the C18 and C11 vesicle solutions, respectively. The solution was then set aside for 15 min to equilibrate, after which the average size of the particles in the solution was measured using Zetasizer Nano ZS90, (Malvern Panalytical Ltd., Malvern, UK). The average size of the population was plotted against the $Mg^{2+}$ ion concentration, so as to estimate $Mg^{2+}$ ion induced fatty acid aggregation. The total lipid concentration was kept at 2 mM for the four C18 based systems. 20 mM lipid solution was used for all the three heterogeneous C11 based systems to prevent concentration induced vesicle aggregation. However, for the homogenous UDA system, the lipid concentration was kept at 60 mM due to its intrinsically high CVC. As for the microscopy analysis, the lipid concentration was kept the same as was used for the DLS experiment. Desired amount of $Mg^{2+}$ ion concentration was obtained by adding different volumes of the $MgCl_2$ stock solution in the lipid solution. Presence of different forms of aggregates, namely, vesicles, lipid crystals and oil droplets, were checked for using DIC microscopy.

### Permeability Assay

In order to determine the permeability of the different membrane systems, calcein leakage assay was used. Calcein is a small polar molecule with an excitation and emission wave length of 495 and 515 nm, respectively, and it gets self-quenched at a high concentration. This property was used to carry out this study by encapsulating calcein above its self-quenching concentration. This was done by rehydrating the dried lipid film with 200 mM bicine buffer pH 8, containing 35 mM of calcein. For all the three heterogeneous C11 based systems, 90 mM of total lipid concentration was used. However for the homogenous UDA system, the lipid concentration was kept at 150 mM due to its relatively high CVC. Interestingly, the pH of the solution dropped after dissolving the dried fatty acid film in the aforementioned mix of buffer and calcein. Therefore, the pH of the solution was readjusted by adding NaOH solution. The solution then went through four freeze-thaw cycles to increase the encapsulation efficiency.

To confirm the encapsulation of calcein in the vesicles, the crude suspension was observed under microscope, using both, fluorescence and DIC, as shown in Figure S10B. Thereafter, the calcein encapsulated vesicle solution was extruded 15 times through a 200 nm size cut-off polycarbonate membrane using Avanti mini extruder (Avanti Polar Lipids Inc., Alabaster, AL, USA) and was loaded on to a size exclusion column (20 cm X 1 cm) packed with Sephadex G-50 fine beads. The column was pre-equilibrated with the mobile phase, which is 200 mM bicine buffer containing just empty lipid vesicles. For each lipid system, the mobile phase contained the same lipid composition and ratio, but at a slightly higher concentration than their CVC, to prevent the lysis of the calcein encapsulated vesicles in the column. Fractions were then collected manually and loaded on to a 96-microwell plate (about 220 µL/well). The fluorescence was measured using a 96-well plate Varioskan Flash multimode reader (Thermo Scientific, Singapore), by exciting the samples at 495 nm and measuring the emitted light at 515 nm. The vesicles with encapsulated calcein eluted in the early fractions (fraction number 11 to 16), and the unencapsulated calcein eluted in the later fractions (fraction number 43 to 57), as shown in figure S10A. The fluorescence was monitored continuously for three hours. After that, 4 µL of Triton 100X was added in each well to rupture the vesicles and release the remaining encapsulated calcein, which led to maximum fluorescence.

The percentage of encapsulation was calculated by using the following equation.

Encapsulation (%) = $100 * (1 - \frac{Ft-F0}{Ff-F0})$

Where, $F0$ is the fluorescence at time zero, $Ft$ is the fluorescence at time t and $Ff$ is the final fluorescence after the addition of Triton 100X.

**Zeta potential measurement of the lipid solutions**

To determine the negative charge density on the vesicles, the lipid solution was extruded 15 times through a 200 nm size cut-off polycarbonate membrane using Avanti mini extruder (Avanti Polar Lipids Inc., Alabaster, AL, USA). For all the four C18 systems, a total of 2 mM lipid solution was used. In case of all the three mixed C11 systems and homogenous UDA system, 20 mM and 60 mM lipid solutions, respectively, were used. The lipid solutions were kept for one hour to equilibrate before the actual measurement was taken. Post equilibration 600 ul of lipid solution was loaded in a cuvette and the zeta potential readings were acquired using a Zetasizer Nano ZS90 (Malvern Panalytical Ltd., Malvern, UK).

**LC-MS analysis of free fatty acid**

**Sample preparation**

In case of C11 based systems, 60 mM and 90 mM lipid concentration was used, for the UDA and the three mixed systems, respectively. This was done to keep the concentration of the fatty acid i.e. UDA constant (60 mM) across all four systems. The lipid solution was prepared in 200 mM bicine buffer of pH 8. Similarly, for the C18 systems, 20 mM and 30 mM lipid concentration was used for the homogenous

OA and the three mixed systems to keep the concentration of the oleic acid constant (20 mM) across all four systems.  The lipid solution were prepared in 100 mM CHES buffer of pH 9. 500 µL of each sample was loaded onto a Vivaspin 2 centrifugal concentrator, with a molecular weight cut-off of 3 kDda and centrifuged at 5,000g for 15 min. Typically, 50 µL of the filtrate was collected. This filtrate was then acidified by adding 5 µl of formic acid. The free fatty acids present in the filtrate was then extracted by adding 400 µL of 2:1 choloroform:methanol solution. This solution also contained myristoleic acid of 10 µM as an internal standard. The extraction was carried out by vortexing the solution rigorously and then spinning it at 3000g for 2 mins. The organic phase was withdrawn carefully and was dried under a stream of nitrogen gas, and subsequently re-dissolved into 400 ul of 2:1 choloroform:methanol. A fraction of this was loaded on to the column (details in the below section).

**Fatty acid quantification**

Separation of fatty acids was carried out using a Luna C18 column from Phenomenex, Torrance, CA, USA (dimensions: 4.6 X 250 mm, 5 nm particle size). The solvent system used for the liquid chromatography part was as follows: Buffer A used was 95:5 of water : methanol (vol/vol) + 0.1% ammonium hydroxide, and Buffer B was 60:35:5 of isopropanol: methanol : water (vol/vol) + 0.1% ammonium hydroxide. A typical LC run's total time was 22 minutes long. The gradient involved an increase in solvent B concentration from 5% to 100% in 4 minutes, followed by an isocratic phase of 100% of solvent B for fifteen minutes. This was followed by an equilibration phase for four minutes with solvent A, all of which was done at a constant flow rate of 0.4 mL/min. The undecylenic acid and oleic acid (from the respective reaction mixtures), and the myristoleic acid internal standard, eluted at 15.66, 15.75 and 15.84 minutes, respectively. The mass spectrometry was carried out on a Sciex X500R QTOF mass spectrometer (MS) fitted with an Exion-LC series UHPLC (Sciex, CA, USA), using Information Dependent Acquisition (IDA) scanning method. All the mass acquisitions were performed using Electron spray ionization (ESI) in the negative mode with the following parameters: turbo spray ion source, medium collision gas, curtain gas = 30 L/min, ion spray voltage = -4500 V (negative mode), at 300 °C. TOF-MS acquisition was done at declustering potential of -80 V, while using -10 V collision energy. The acquired data was analyzed using the Sciex OS software (Sciex, CA, USA; University of Florida, FL, USA). The presence of a specific species was confirmed by the presence of precursor mass within 3 ppm error range. The fatty acid was quantified by taking the ratio of the area under the corresponding fatty acid peak, with respect to the area of internal standard peak.

**Membrane stability under Multiple Selection Pressures (MSPs)**

To evaluate the stability of protocell membranes as a function of their composition under multiple selection pressures, C11 based membrane systems were used. Three environmental selection pressures, i.e. A) vesicle formation in alkaline pH regime. B) vesicle stability in dilution regimes, and C) vesicle stability in the presence of $Mg^{2+}$ ions, were applied to the systems in a sequential manner. In order to understand if there was any effect coming from the sequence of the applied selection on the fitness of the vesicles, the selection pressures were applied in all possible combinations. Therefore, a total of six different sequential combinations were investigated. All the four C11 based systems were prepared in 200 mM bicine buffer at pH 8 with the lipid concentration kept at 60 mM. Typically, 300 µL of each vesicle suspension was taken in a centrifuge tube. In one of the MSPs sequence combinations tested, first the lipid solution was diluted with 200 mM bicine buffer of pH 8., in order to check the stability of the vesicles on dilution. In the next step, the pH was increased by adding desired volume of 3M NaOH solution. It was added to all the lipid solutions to bring the final pH to 10. In the final step, the stability was checked in the presence of $Mg^{2+}$ ions by adding desired amount of $MgCl_2$ solution to reach a concentration of 14 mM $Mg^{2+}$. After the application of each aforesaid selection pressure, the lipid suspension was observed under microscope at 40X magnification to check for the presence of vesicle and unordered aggregates. In the same manner, the other five MSP combinations were also tested and the membrane systems were evaluated using microscopic analysis.

**Table TS1:** Summary of aggregation-inducing $Mg^{2+}$ ion concentration ($Mg^{2+}{}_{AIC}$) for all different C18 and C11 based systems using two different assays: Columns 2 and 3 provide a comparison of the difference in the $Mg^{2+}{}_{AIC}$ estimation using DLS and microscopy assay, respectively. The fatty acid to overall derivative molar ratio was kept to 2:1. UDA, undecylenic acid; UDG, glyceryl 1-undecylenate; UDOH, undecylenyl alcohol; OA, oleic acid; GMO, glycerol 1-monooleate; OOH, oleyl alcohol.

| | Mg2+ ion (in mM) induced aggregation formation | |
|---|---|---|
| System used | DLS analysis | Microscopy |
| UDA | 3 | 3 |
| UDA:UDOH | 8 | 8 |
| UDA:UDG | 16 | 12 |
| UDA:UDG:UDOH | 14 | 12 |
| OA | 3.5 | 2 |
| OA:OOH | 6 | 6 |
| OA:GMO | 5 | 4.5 |
| OA:GMO:OOH | 6 | 6 |

**Figures S1 to S17**

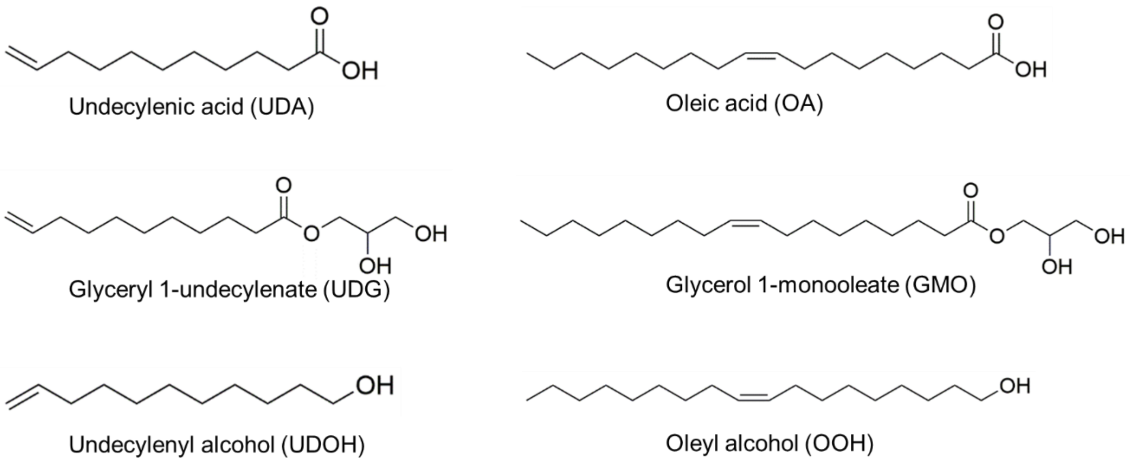


**Figure S1:** Structures of different amphiphiles, i.e. fatty acids (C11 and C18) and their derivatives used in the present study.

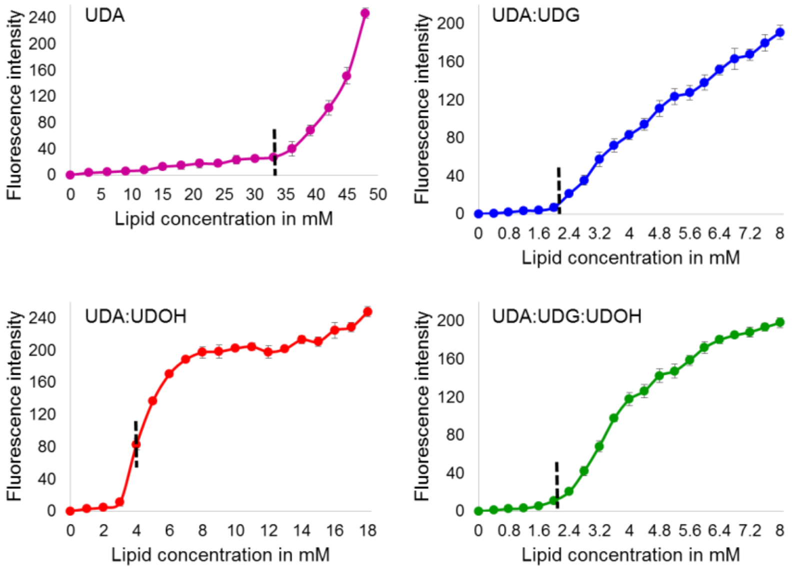


**Figure S2:** CVC estimation of different C11 membrane systems by fluorescence assay. The increase in fluorescence is plotted as a function of lipid concentration. The inflection point, which is a read out of the CVC of the system, is represented with the black dashed line. The fatty acid to overall derivative molar ratio was kept to

2:1. UDA, undecylenic acid; UDG, glyceryl 1-undecylenate; UDOH, undecylenyl alcohol. n = 3; error bars represent standard deviation (s.d.).

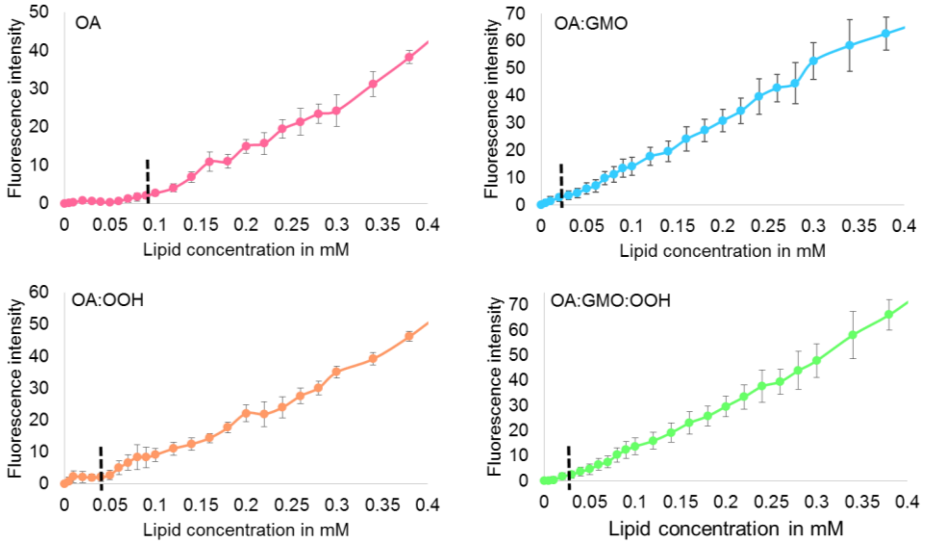


**Figure S3:** CVC estimation of the different C18 membrane systems by fluorescence assay. The increase in fluorescence is plotted as a function of lipid concentration. The inflection point, which is a read out of the CVC of the system, is represented with the black dashed line. The fatty acid to overall derivative molar ratio was kept to 2:1. OA, oleic acid; GMO, glycerol 1-monooleate; OOH, oleyl alcohol. n = 3; error bars represent standard deviation (s.d.).

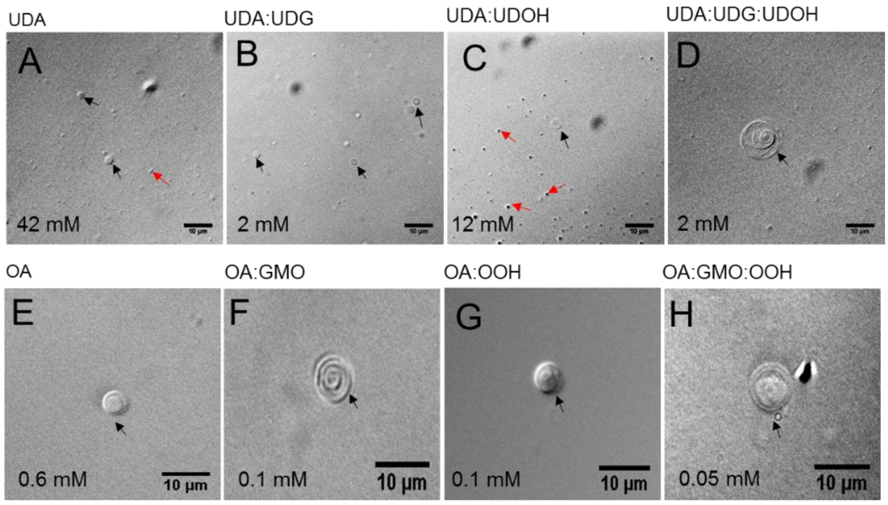


**Figure S4:** CVC estimation using microscopic analysis. Panels A to D and E to F show the microscopic analysis of the different C11 and C18 based systems, respectively, at their CVC. A) 40 mM homogenous UDA, B) 2 mM of UDA:UDG, C) 12 mM of UDA:UDOH, D) 2 mM of the tertiary UDA:UDG:UDOH system, E) 0.6 mM of OA, F) 0.1 mM of OA:GMO, G) 0.1 mM of 0.1 mM OA:UDOH, and H) 0.05 mM of OA:GMO:OOH systems. Below the aforesaid concentrations, vesicles were not observed under 40X magnification. The black and red arrows indicate vesicles and aggregates, respectively. Scale bar in all the images is 10 microns.

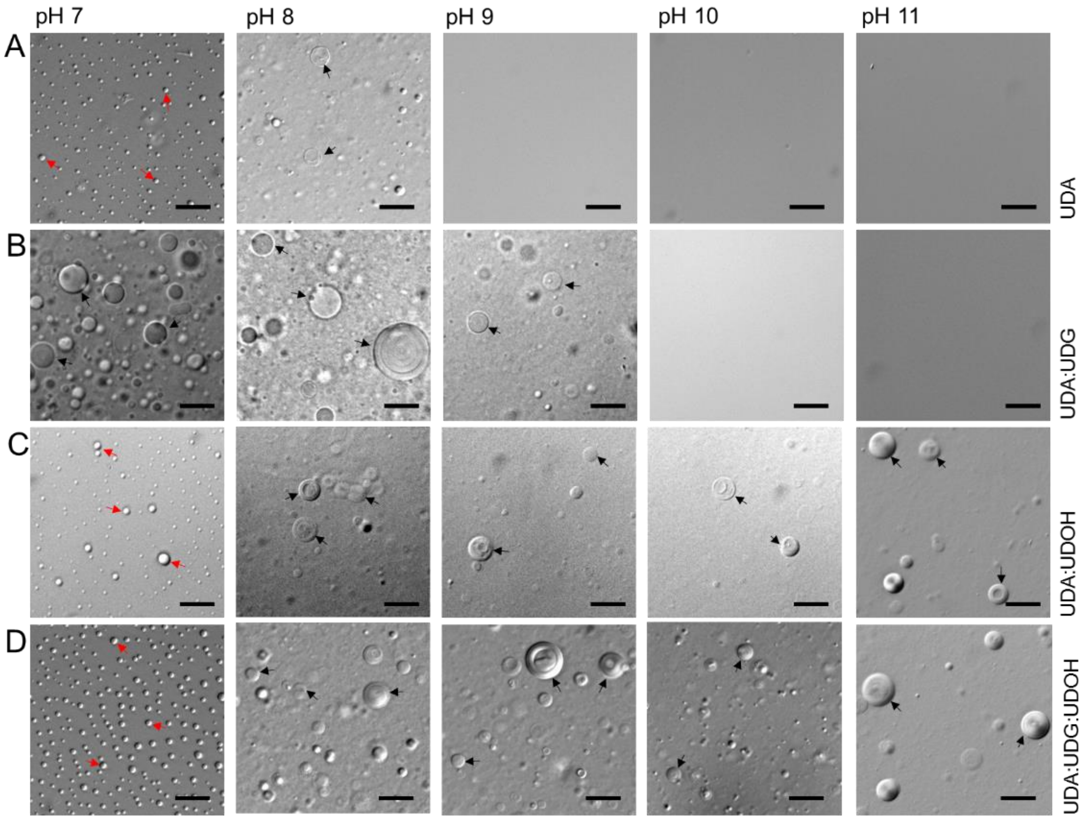


**Figure S5:** Microscopic analysis of C11 membrane systems. These images demonstrate the formation of different higher order assemblies (e.g. vesicles, oil droplets etc.), depending on the pH of the surrounding environment. Panels A to D show the four different C11-based systems. The black and red arrows indicate vesicles and aggregates, respectively. The scale bar in all the images is 10 microns.

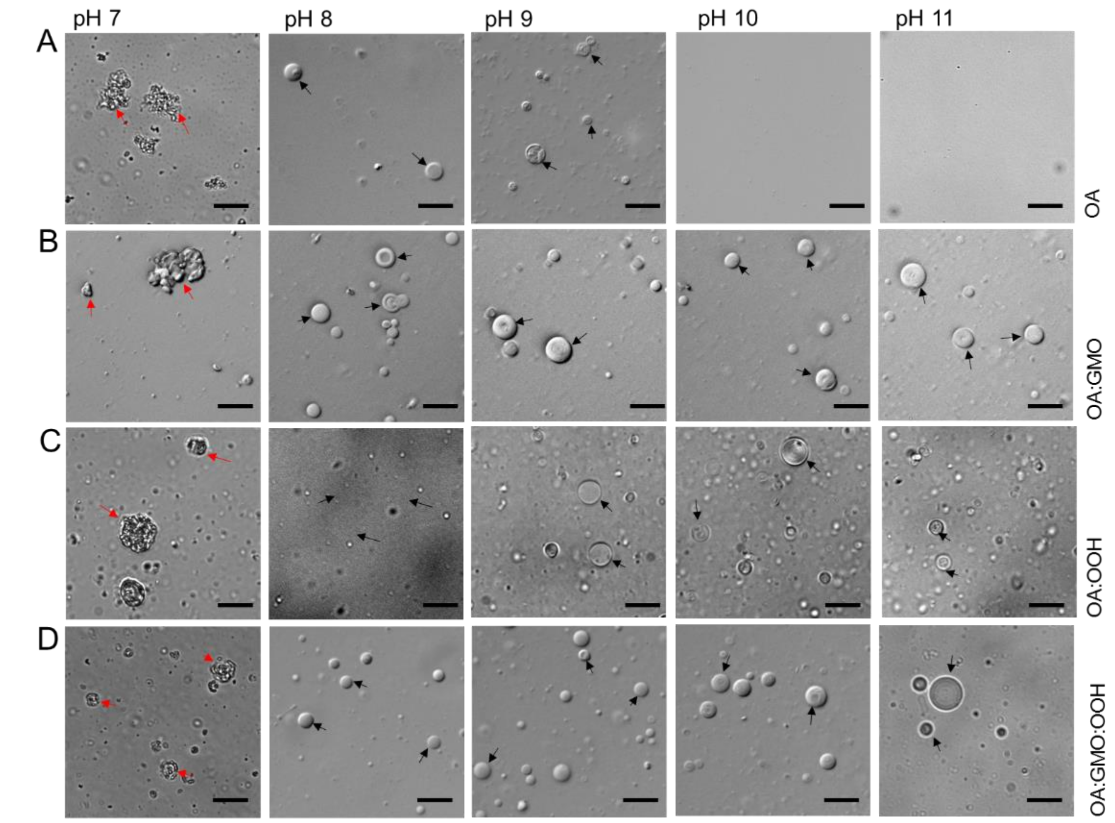


**Figure S6:** Microscopic analysis of C11 membrane systems. These images demonstrate the formation of different higher order assemblies (e.g. vesicles, oil droplets etc.), depending on the pH of the surrounding environment. Panels A to D show the four different C18-based systems. The black and red arrows indicate vesicles and aggregates, respectively. The scale bar in all the images is 10 microns.

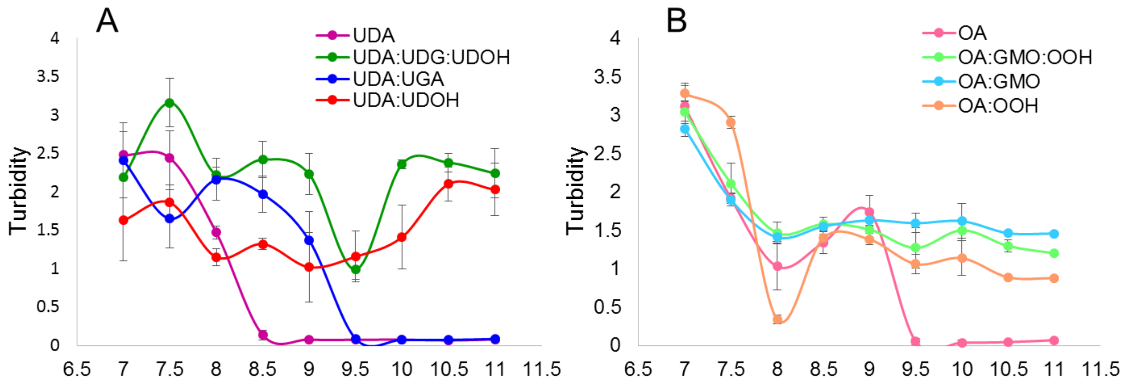


**Figure S7:** Turbidity measurements of different C18 and C11 membrane systems at different pH. The turbidity of the systems at 400 nm is plotted as a function of the pH. Panel A and B represent the different C11 and C18 systems, respectively. Decrease in turbidity indicates the formation of micelles in the system, which cannot scatter light at 400 nm.

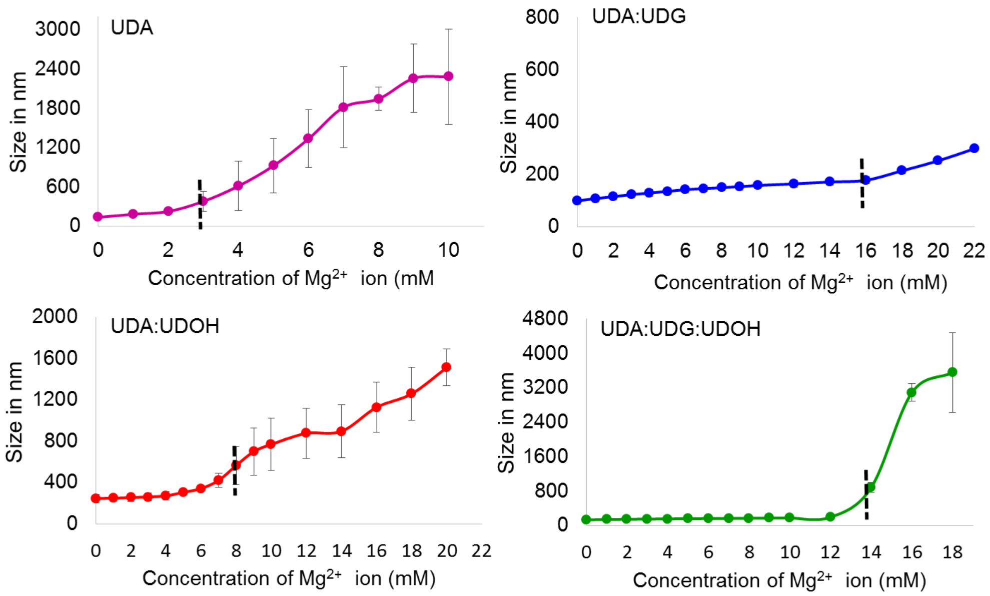


**Figure S8:** DLS measurements of C11 membrane systems to determine the $Mg^{2+}$ ion induced aggregate formation concentration. The particle diameter (in nm) is plotted as a function of the added $Mg^{2+}$ ion concentration. The vertical black dashed line indicates the $Mg^{2+}$ ion induced aggregation formation concentration ($Mg^{2+}_{AIC}$). n = 3; error bars represent standard deviation (s.d.).

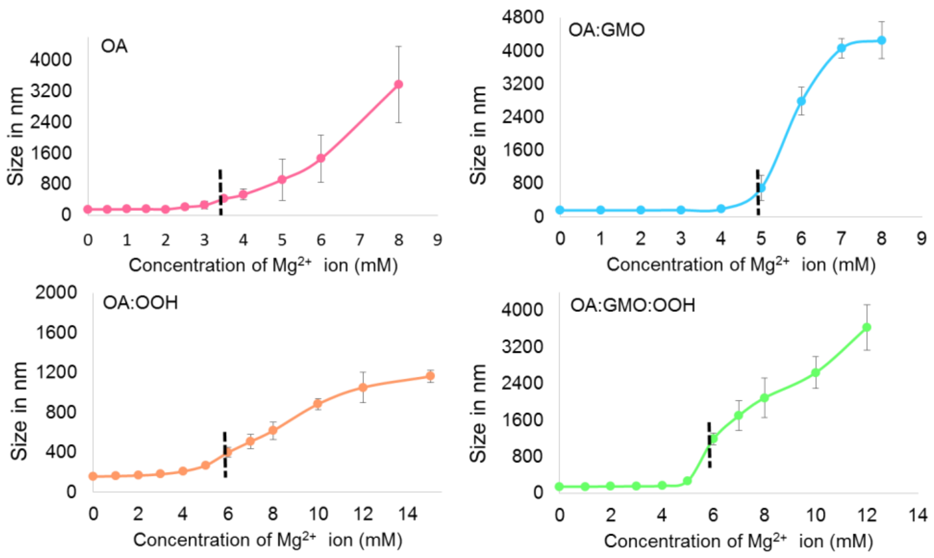

**Figure S9:** DLS measurements of the C18 membrane systems to determine $Mg^{2+}$ ion induced aggregate formation concentration. The particle diameter (in nm) is plotted as a function of the added $Mg^{2+}$ ion concentration. The vertical black dashed line indicates the $Mg^{2+}$ ion induce aggregation formation concentration ($Mg^{2+}_{AIC}$). n = 3; error bars represent standard deviation (s.d.).

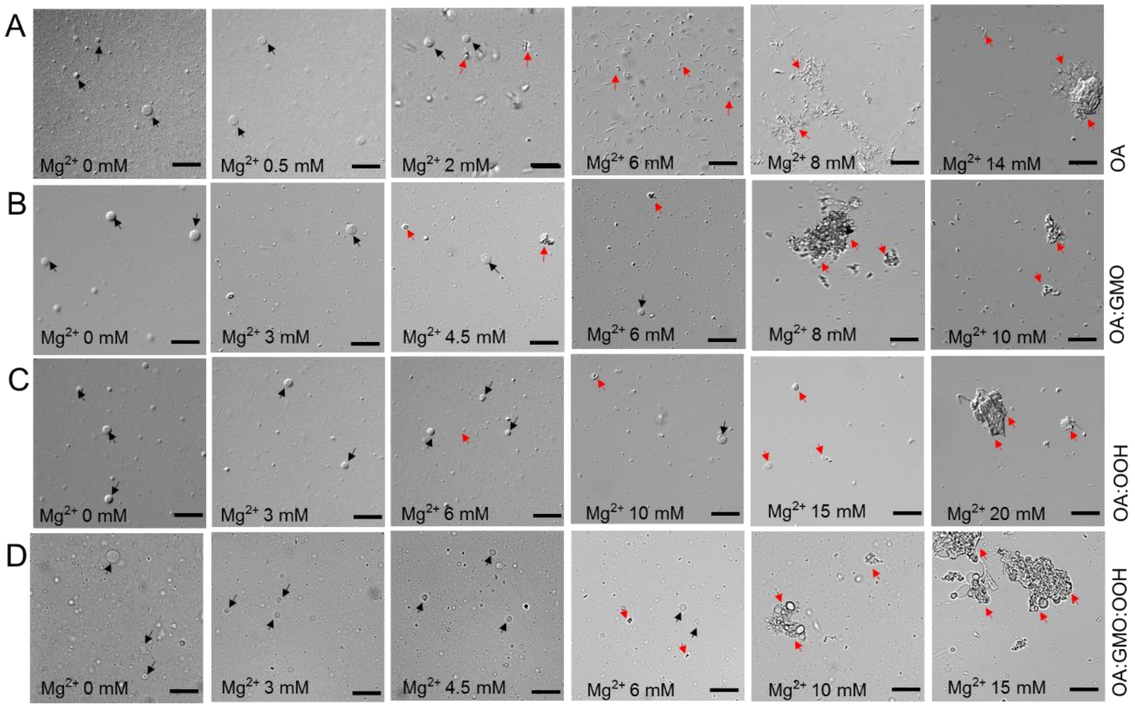


**Figure S10:** $Mg^{2+}$ ion induced vesicle and aggregate forming properties of all the four C18 based membrane systems (Panels A to D). From left to right, $Mg^{2+}$ ion concentration was increased gradually by keeping the lipid concentration constant. In terms of cation sensitivity among the four systems, the following order is observed: OA > OA:GMO > OA:GMO:OOH = OA:OOH. The black and red arrows indicate vesicles and aggregates (fatty acid crystals and droplets), respectively. The scale bar in all images is 20 microns.

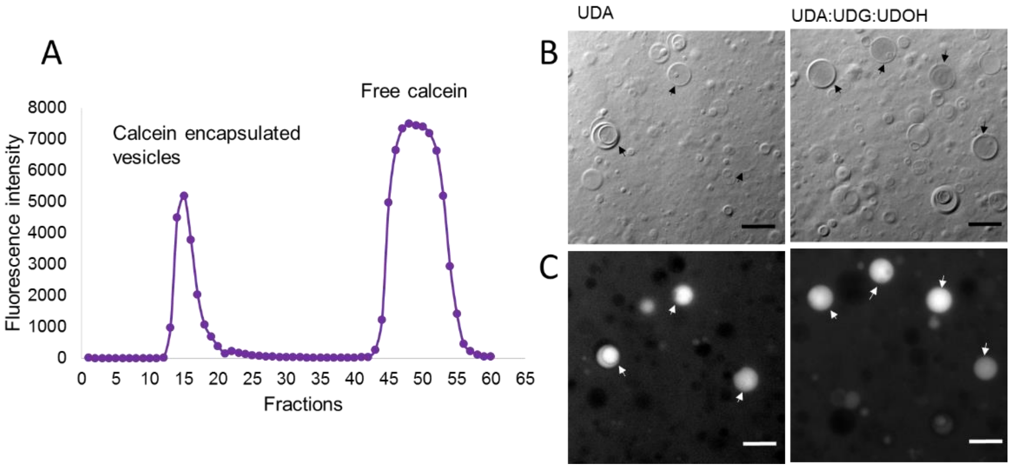


**Figure S11:** S11A shows the size exclusion column chromatography profile, which was used to separate vesicles with encapsulated calcein, from the unencapsulated calcein. Fluorescence at 518 nm was plotted against the fraction number. The unencapsulated calcein comes out in later fractions whereas the vesicles with encapsulated calcein elute out in the earlier fractions. S11B shows the epifluorescence micrographs of calcein encapsulated vesicles from two C11 based membrane systems. The images on the top represent the Differential Interference Contrast (DIC) images, while the lower two images are of the fluorescence images of the same field of view. The scale bar in all the images is 10 microns.

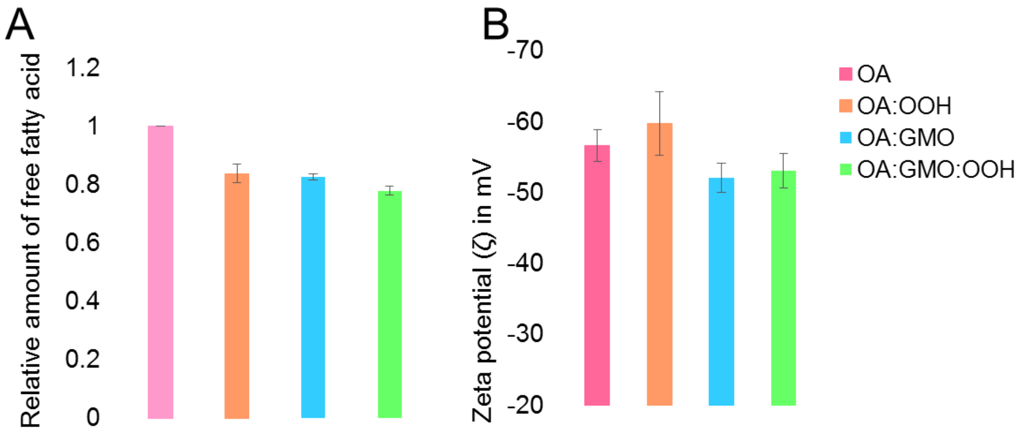


**Figure S12:** S12A represents the relative amount of free oleic acid molecules (C18) present in the solution as a function of membrane composition in all four C18 membrane systems. n = 6; error bars represent standard deviation (s.d.). The difference between the means for homogenous OA and the other three heterogeneous systems is significant based on student t test with a *p*-value < 0.05 (using a one-tailed test). S12B represents the zeta potential measurements of the C18 systems as a function of their composition. n = 5; error bars represent standard deviation (s.d.). The difference between the means of the OA system and that of theOA:GMO/ OA:GMO:OOH systems is significant based on student t test; with a *p*-< 0.05. The difference between the means obtained for the homogenous OA and the binary OA:OOH is not significant based on student t test: *p*-value > 0.05 (using a one-tailed test).

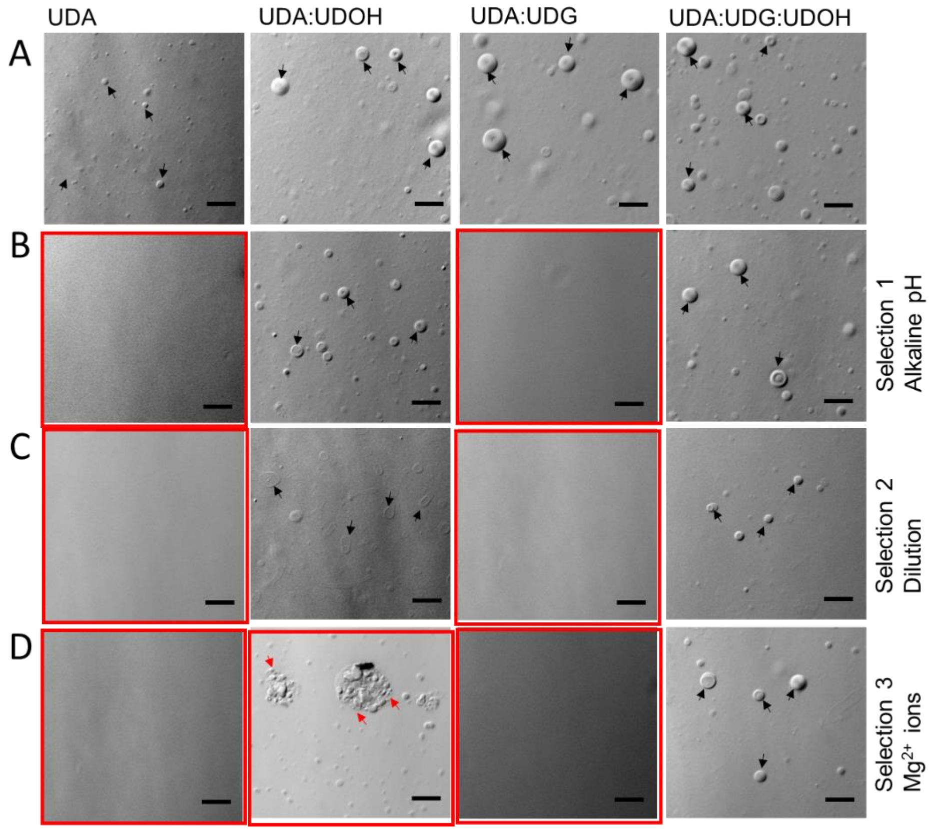


**Figure S13:** Vesicle stability as a function of their composition under multiple selection pressures (MSPs), when applied consecutively. Panels B to D represent the different pebiotically relevant selection conditions. A) All four C11 systems at a concentration of 60 mM at pH 8. B) Stability at alkaline pH as a selection pressure. All systems comprised of 60 mM of lipid concentration, with the pH of the system now adjusted to 10. C) Dilution regime as selection pressure in which all the four systems were diluted to concentration of 20 mM lipid concentration at pH 10. D) Stability in the presence of $Mg^{2+}$ ions as selection pressure. $Mg^{2+}$ ions were added in all the systems at a concentration of 14 mM. The lipid concentration is 20 mM in all the systems, which are at pH10. The red boxes indicate conditions where the vesicles are absent. The black and red arrows indicate vesicles and aggregates (fatty acid crystals and droplets), respectively. The scale bar in all the images is 10 microns.

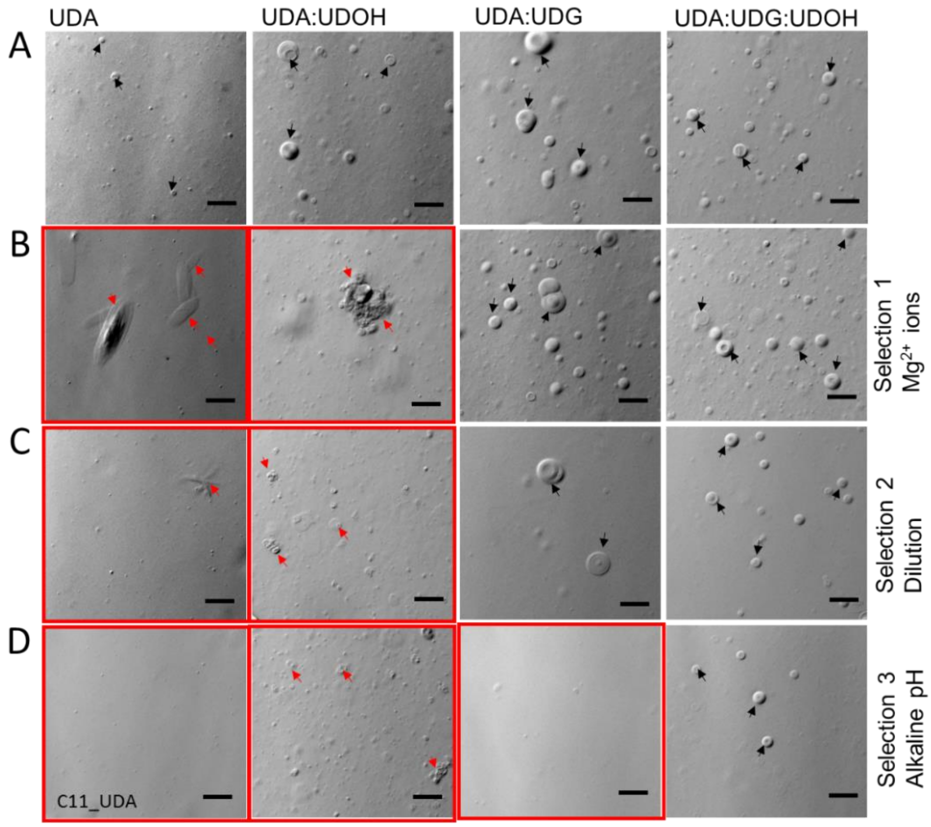


**Figure S14:** Vesicle stability as a function of their composition under multiple selection pressures (MSPs), when applied consecutively. Panels B to D represent the different pebiotically relevant selection conditions. A) All four C11 systems at a concentration of 60 mM at pH 8. B) Stability in the presence of $Mg^{2+}$ ions as selection pressure. $Mg^{2+}$ ions were added in all the systems at a concentration of 14 mM. The lipid concentration is 60 mM in all the systems, which are at pH8. C) Dilution regime as selection pressure in which all the four systems were diluted to concentration of 20 mM lipid concentration at pH 8, in presence of 14 mM $Mg^{2+}$ ions. D) Stability at alkaline pH as a selection pressure. All systems comprised of 20 mM of lipid concentration containing 14 mM $Mg^{2+}$ ions, with the pH of the system now adjusted to 10. The red boxes indicate conditions where the vesicles are absent. The black and red arrows indicate vesicles and aggregates (fatty acid crystals and droplets), respectively. The scale bar in all the images is 10 microns.

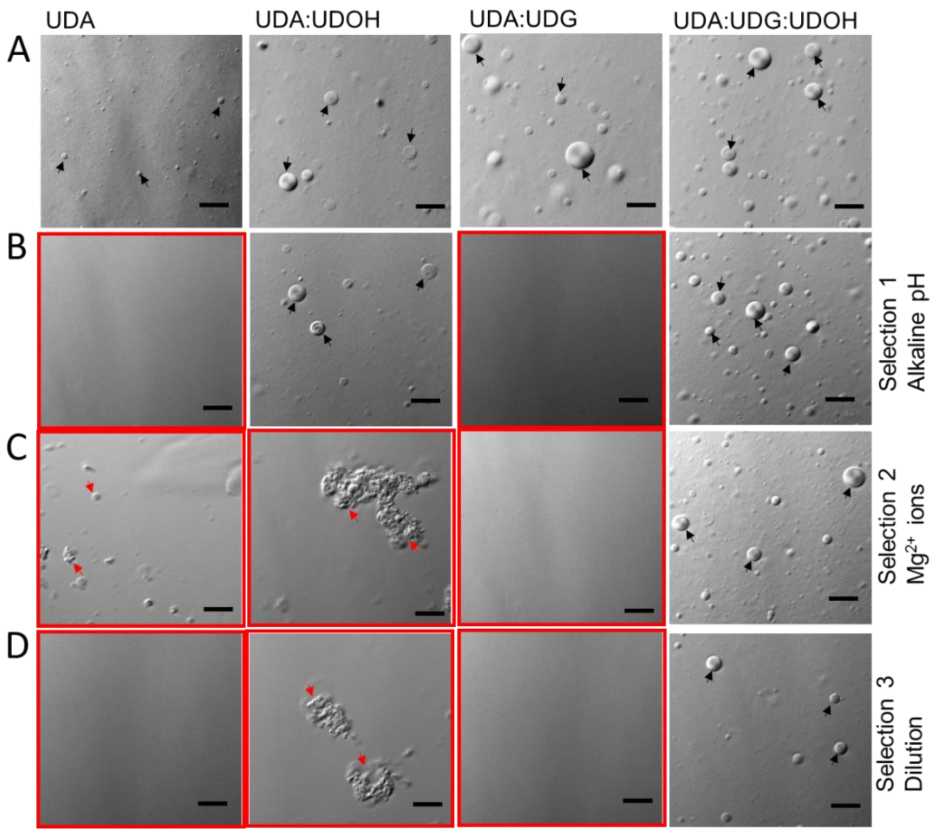


**Figure S15:** Vesicle stability as a function of their composition under multiple selection pressures (MSPs), when applied consecutively. Panels B to D represent the different pebiotically relevant selection conditions. A) All four C11 systems at a concentration of 60 mM at pH 8. B) Stability at alkaline pH as a selection pressure. All systems comprised of 60 mM of lipid concentration, with the pH of the system now adjusted to 10. C) Stability in the presence of $Mg^{2+}$ ions as selection pressure. $Mg^{2+}$ ions were added in all the systems at a concentration of 14 mM. The lipid concentration is 60 mM in all the systems, which are at pH 10. D) Dilution regime as selection pressure in which all the four systems were diluted to concentration of 20 mM lipid concentration at pH 10 in presence of 14 mM $Mg^{2+}$ ions. The red boxes indicate conditions where the vesicles are absent. The black and red arrows indicate vesicles and aggregates (fatty acid crystals and droplets), respectively. The scale bar in all the images is 10 microns.

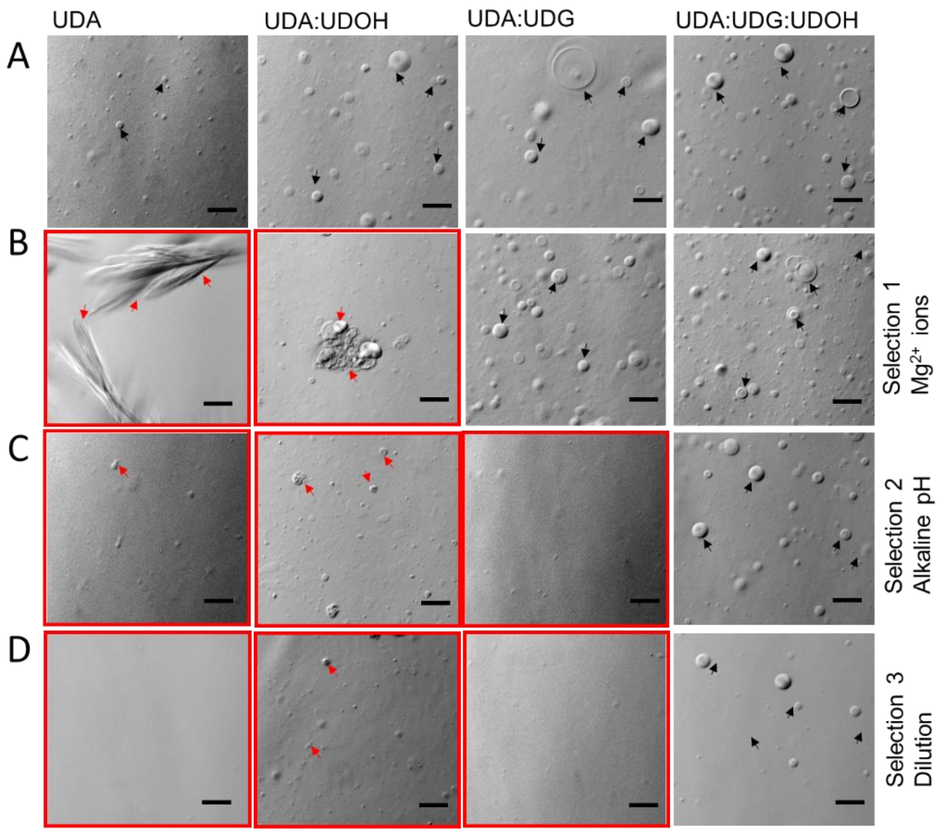


**Figure S16:** Vesicle stability as a function of their composition under multiple selection pressures (MSPs), when applied consecutively. Panels B to D represent the different pebiotically relevant selection conditions. A) All four C11 systems at a concentration of 60 mM at pH 8. B) Stability in the presence of $Mg^{2+}$ ions as selection pressure. $Mg^{2+}$ ions were added in all the systems at a concentration of 14 mM. The lipid concentration is 60 mM in all the systems, which are at pH8. C) Stability at alkaline pH as a selection pressure. All systems comprised of 60 mM of lipid concentration and 14 mM $Mg^{2+}$ ions, with the pH of the system now adjusted to 10. D) Dilution regime as selection pressure in which all the four systems were diluted to concentration of 20 mM lipid concentration at pH 10, in presence of 14 mM $Mg^{2+}$ ions. The red boxes indicate conditions where the vesicles are absent. The black and red arrows indicate vesicles and aggregates (fatty acid crystals and droplets), respectively. The scale bar in all the images is 10 microns.

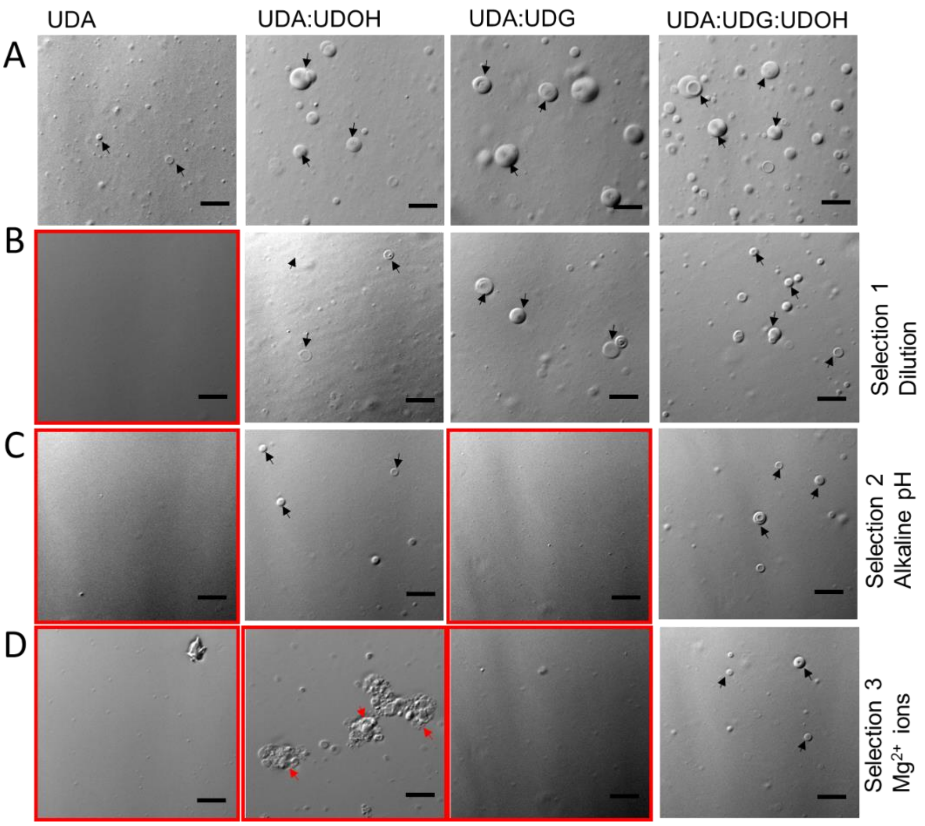


**Figure S17:** Vesicle stability as a function of their composition under multiple selection pressures (MSPs), when applied consecutively. Panels B to D represent the different pebiotically relevant selection conditions. A) All four C11 systems at a concentration of 60 mM at pH 8. B) Dilution regime as selection pressure in which all the four systems were diluted to concentration of 20 mM lipid concentration at pH 8. C) Stability at alkaline pH as a selection pressure. All systems comprised of 20 mM of lipid concentration, with the pH of the system now adjusted to 10. D) Stability in the presence of $Mg^{2+}$ ions as selection pressure. $Mg^{2+}$ ions were added in all the systems at a concentration of 14 mM. The lipid concentration is 20 mM in all the systems, which are at pH10. The red boxes indicate conditions where the vesicles are absent. The black and red arrows indicate vesicles and aggregates (fatty acid crystals and droplets), respectively. The scale bar in all the images is 10 microns.
